# Protein Posttranslational Signatures Identified in COVID-19 Patient Plasma

**DOI:** 10.1101/2021.12.15.472822

**Authors:** Pavan Vedula, Hsin-Yao Tang, The UPenn COVID Processing Unit, David W. Speicher, Anna Kashina

## Abstract

Severe acute respiratory syndrome coronavirus-2 (SARS-CoV-2) is a highly contagious virus of the coronavirus family that causes coronavirus disease-19 (COVID-19) in humans and a number of animal species. COVID-19 has rapidly propagated in the world in the past 2 years, causing a global pandemic. Here, we performed proteomic analysis of plasma samples from COVID-19 patients compared to healthy control donors in an exploratory study to gain insights into protein-level changes in the patients caused by SARS-CoV-2 infection and to identify potential proteomic and posttranslational signatures of this disease. Our results suggest a global change in protein processing and regulation that occurs in response to SARS-CoV-2, and the existence of a posttranslational COVID-19 signature that includes an elevation in threonine phosphorylation, a change in glycosylation, and a decrease in arginylation, an emerging posttranslational modification not previously implicated in infectious disease. This study provides a resource for COVID-19 researchers and, longer term, will inform our understanding of this disease and its treatment.

**Key Points:**

1. Plasma from COVID-19 patients exhibits prominent protein- and peptide-level changes
2. Proteins from COVID-19 patient plasma exhibit prominent changes in several key posttranslational modifications

## Introduction

Severe acute respiratory syndrome coronavirus-2 (SARS-CoV-2) is a respiratory virus of the coronavirus family that causes coronavirus disease-19 (COVID-19) in humans and a number of animal species ^1^. COVID-19 has rapidly propagated worldwide in the past 2 years, causing a global pandemic (see, e.g., ^2^ for a recent review). This highly contagious disease causes respiratory symptoms that range from mild to severe, and is associated with a number of other serious health implications, including lung inflammation and damage, thrombosis, stroke, renal failure, neurological disorders, and others^3-8^. This list continues to grow, and despite extensive research in the past year and a half, full understanding of COVID-19 mechanisms of action and health consequences has not yet been achieved.

Here, we performed proteomic analysis of plasma samples from COVID-19 patients with symptoms severe enough to require hospitalization compared to healthy control donors, in an attempt to gain insights into protein-level changes in the patients caused by SARS-CoV-2 infection and to identify potential proteomic and posttranslational signatures of this disease. Our analysis revealed a number of changes in protein and peptide composition of the COVID-19 patients’ plasma samples. Furthermore, global analysis of posttranslational modifications (PTMs) in these samples showed a striking change in several key physiological PTMs, including phosphorylation, glycosylation, citrullination, and arginylation, which exhibited differential up- and down-regulation in COVID-19 patients compared to controls and, in the case of arginylation and phosphorylation, modified different repertoire of sites on a limited number of target proteins. These patterns suggest a global change in protein processing and regulation that occurs in response to SARS-CoV-2, and the existence of a posttranslational COVID-19 “code”. Deciphering this code may advance our understanding of disease progression and long-term implications, as well as potentially inform novel strategies of COVID-19 diagnostics and treatment.

## Results and Discussion

### COVID-19 patients exhibit prominent changes in their plasma peptidomes

To address potential protein and peptide changes in the plasma associated with COVID-19, we obtained plasma samples from 6 COVID patients with severe disease that required hospitalization and 7 similarly drawn control samples from healthy donors collected independently within the same time frame (Table S1).

To analyze plasma peptidomes, we used size exclusion under denaturing conditions followed by C18 reverse phase cleanup to isolate plasma peptides followed by LC-MS/MS without proteolysis treatment. LC-MS/MS data were searched against the human protein database using a no-enzyme specificity so that peptides naturally occurring or produced by in vivo proteolysis could be identified. The final search results were filtered by p-value (<0.05) and fold change (2-fold and above) to define 180 peptides that showed significant differences in abundance between patients and controls. The significantly changed peptides are shown in Table S2, and the list of identified peptides that did not meet these statistical criteria and were not used in the final analysis is shown in Table S3.

No known regulatory peptides were identified, likely due to their lower abundance. All the significant peptides identified constituted proteolytic fragments predominantly from abundant plasma proteins, that were apparently produced by proteolytic events that accompany immune response, cell migration and adhesion, and other physiological processes ^9^. Interestingly, when the total number of identified peptides were compared (Tables S2 and S3) substantially fewer peptides were identified overall in the COVID-19 samples compared to control (Fig. S1), in seeming contrast to the fact that SARS-CoV-2 infection is associated with increased proteolysis^10-14^.

To further analyze COVID-19-dependent peptidomics trends, we used our significantly changed peptide list (Table S2) and grouped the identified peptides by their parent proteins. For each given protein we plotted combined intensities of all significantly changed peptides (Fig. S2), and separately plotted all the combined intensities grouped by protein for ten most abundant hits showing overall increase in control (Fig. 1A) and COVID-19 (Fig. 1B).

**Figure 1.**
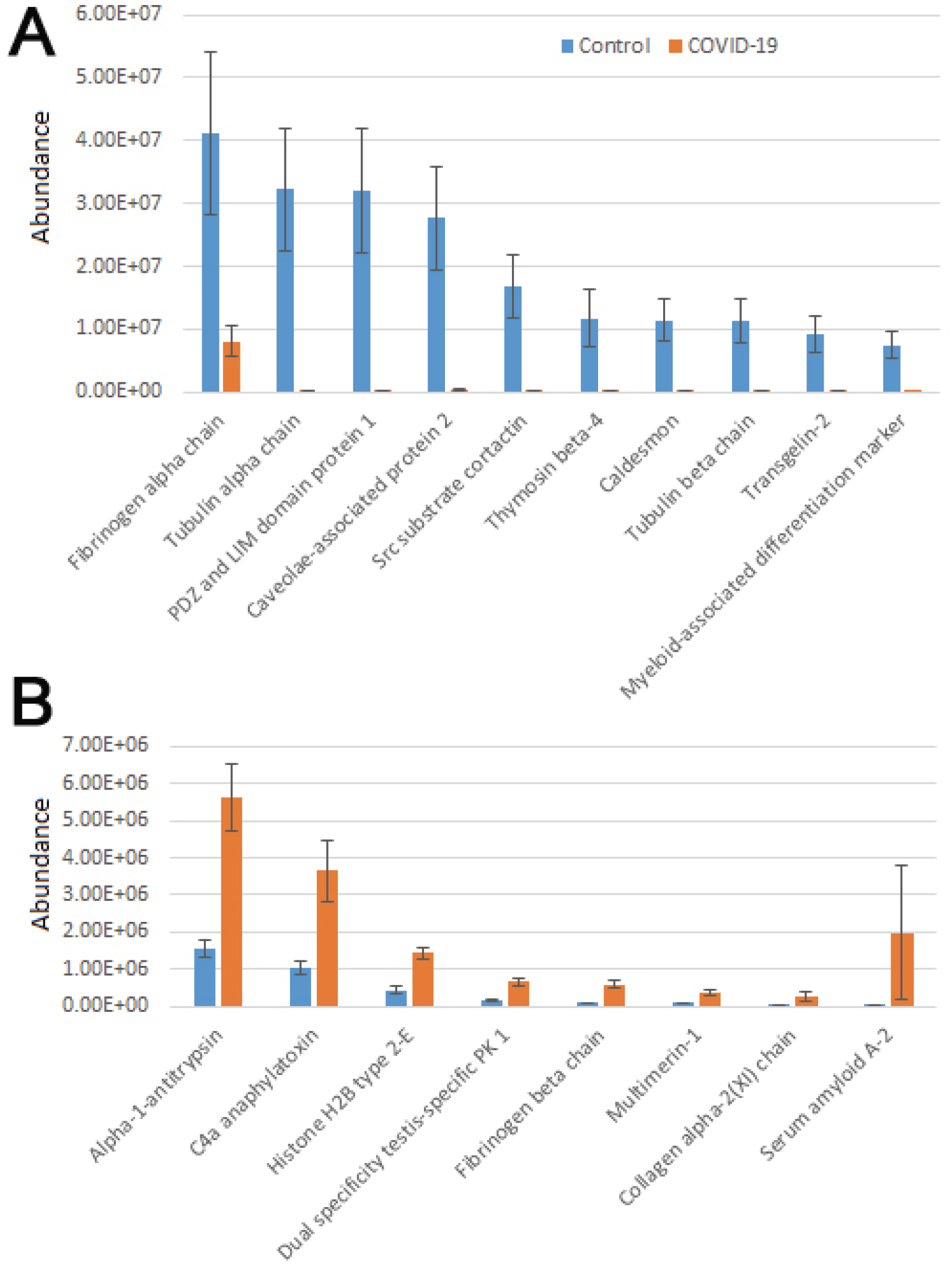
Plasma peptides from COVID-19 patients exhibits prominent changes compared to control. Combined total intensities of all significantly changed peptides for each parent protein listed on the X axis. A and B show top 10 most abundant peptide groups in control (A) and COVID-19 (B). See Fig. S2 for the full list of proteins with significanty changed peptides. Error bars represent SEM (n=7 for control, 6 for COVID-19).

Most of the combined peptide intensities for each protein were substantially higher in the control samples compared to COVID-19, consistent with the fact that control plasma contained more peptides overall (Fig. S1). However, peptides derived from a small group of proteins showed significant elevation in COVID-19 plasma (Fig. 1B). The most abundant of these, Alpha-1-antitrypsin, C4a anaphylatoxin, and Serum amyloid A-2, have known functional association with disease pathology. Alpha-1-antitrypsin is a protease inhibitor, involved in regulation of plasma proteolysis. C4 anaphylatoxin and Serum amyloid A-2 are involved in immune response. Both of these processes are highly relevant to the COVID-19 disease ^3,15-17^, and thus increased abundance of significantly changed peptides for these proteins in COVID-19 patients may indicate a direct association between these proteins’ proteolysis and SARS-CoV-2 infection. In contrast, peptide groups showing higher levels in control belong to normal proteins expected to be proteolyzed in the blood due to normal organismal functions. We speculate that their higher apparent abundance in control compared to COVID-19 samples in our analysis is the result of their decrease in COVID-19.

An additional interesting observation concerns alpha-fibrinogen, which had the largest number of significantly changing peptides (Table S2). Even though the combined intensities of significantly changed fibrinogen-derived peptides was much higher in control compared to COVID-19 (Fig. 1A and S2), individual peptides derived from alpha-fibrinogen showed different trends, with some peptides elevated in COVID-19 rather than control samples (Fig. 2A and S3). The peptides with the highest abundance were still more prevalent in control (Fig. 2A, left), however out of the 17 peptides showing significant change, 8 were more abundant in the COVID-19 samples (Fig. 2A, right). A somewhat similar pattern was observed with serglycin: only two peptides were identified, but one of these was much more abundant in control, while the other showed the opposite trend (Fig. 2B). While each change was significant, added together, the intensities of these peptides evened out and showed no change in the serglycin protein group shown in Fig. S2. This observation suggests that fibrinogen and serglycin undergo different proteolytic events in normal physiology and during SARS-CoV-2 infection. Fibrinogen, plays a key role in blood clotting ^18^, a process shown to be impacted in COVID-19 patients ^19^; serglycin is key to the biology of the blood cells^20^. Altered proteolytic patterns of these proteins in COVID-19 may prove to be a potentially interesting biomarker in future studies.

**Figure 2.**
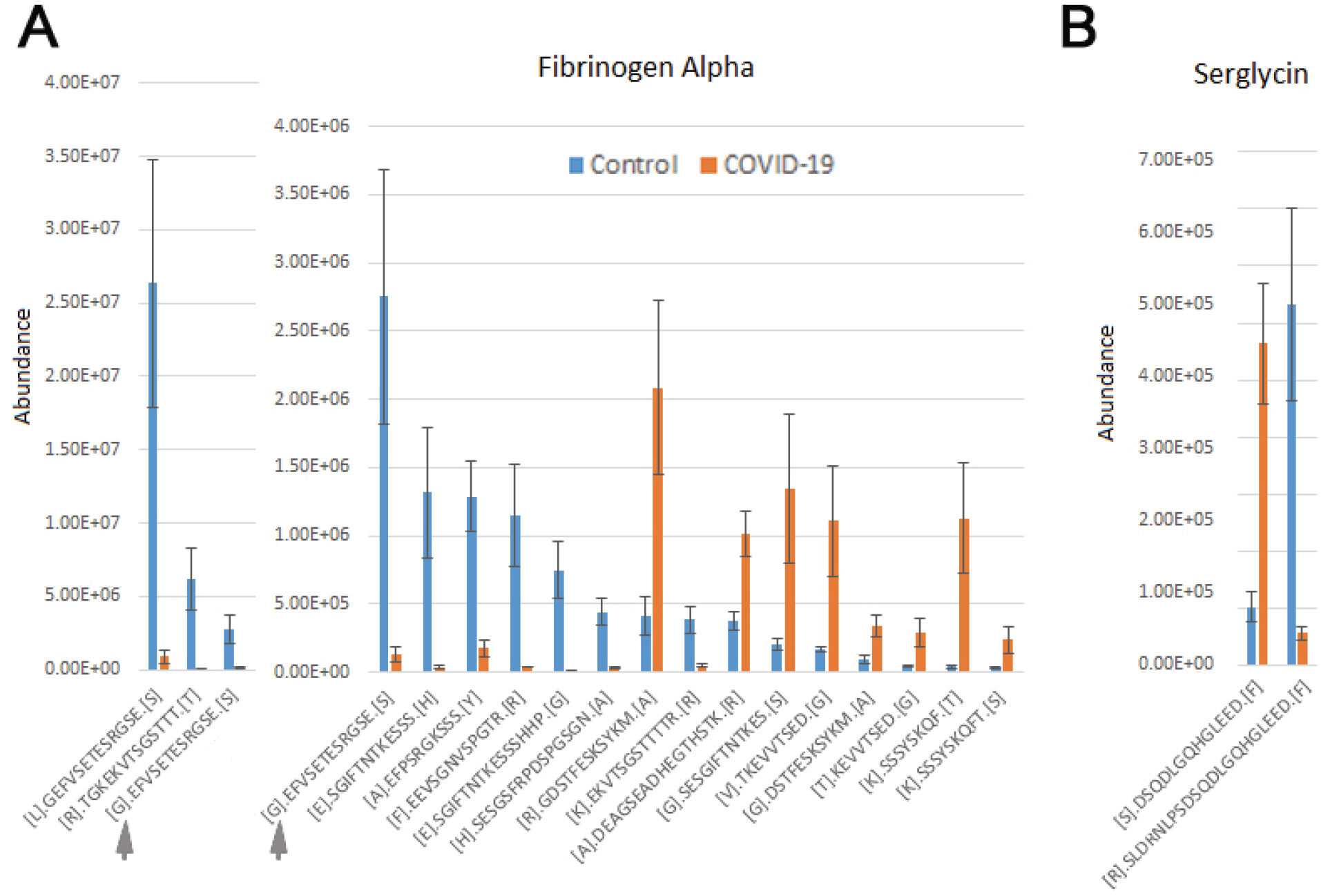
Fibrinogen- and serglycin-derived peptides exhibit differential abundance changes between COVID-19 and control, suggesting different proteolytic patterns in response to Sars-CoV-2 infection. Normalized intensities of the most abundant individual peptides in control and COVID-19 plasma samples are shown for fibrinogen (A) and serglycin (B). Peptides in A are plotted in two charts on two different scales, including one overlapping peptide is shown in both charts for scale ([G].EFVSETESRGSE.[S] indicated with a gray arrow in both charts). Error bars represent SEM (n=7 for control, 6 for COVID-19).

### COVID-19 patient plasma exhibits prominent changes in the global proteome

Next, we analyzed plasma samples by shotgun proteomics. For this, IgG/albumin-depleted plasma samples were loaded onto SDS PAGE and run ∼0.5 cm into the gel, followed by Coomassie Blue staining and excision of the entire protein-containing gel zone, which was then subjected to in-gel digestion with trypsin and analyzed by LC-MS/MS. Differences between COVID-19 and control samples were considered as high confidence significant changes if they exhibited a greater than 2-fold increase or decrease and q-value less than 0.05. Proteins that passed these criteria and were used for further analysis are listed in Table S4. The remaining identified proteins are listed in Table S5.

Interestingly, only 12 proteins were significantly increased in COVID-19 plasma while 35 proteins were decreased in these samples compared with controls (Fig. 3 and S3). When total iBAQ was used as a rough metric of relative abundance across proteins, only three of the 8 most abundant proteins were increased in COVID-19 plasma (Fig. 3). One of these proteins, serum amyloid A-2, also showed up in our peptidomics dataset (Fig. 1), where peptides derived from this protein were elevated to a similar extent in COVID-19 compared to control. This suggests that serum amyloid A-2 is both upregulated and more heavily proteolyzed in COVID-19. The other two proteins increased in COVID-19 plasma were serum amyloid A-1 and C-reactive protein, which are known to be elevated in the plasma in response to inflammation, and thus their increased levels observed in our dataset is fully consistent with known COVID-19 effects. In contrast, proteins showing decreased levels in COVID-19 compared to control (Fig. 3, S3, and Table S3) are mostly related to normal physiological functions, including anti-inflammatory response (apolipoprotein A-IV and C-III) and overall protective functions (serum paraoxonase), hormone and vitamin transport (retinol-binding protein and transthyretin). It is possible that their elevated levels in control versus patient plasma actually reflect down-regulation or depletion of these normal proteins upon SARS-CoV-2 infection that ultimately contribute to disease pathology.

**Figure 3.**
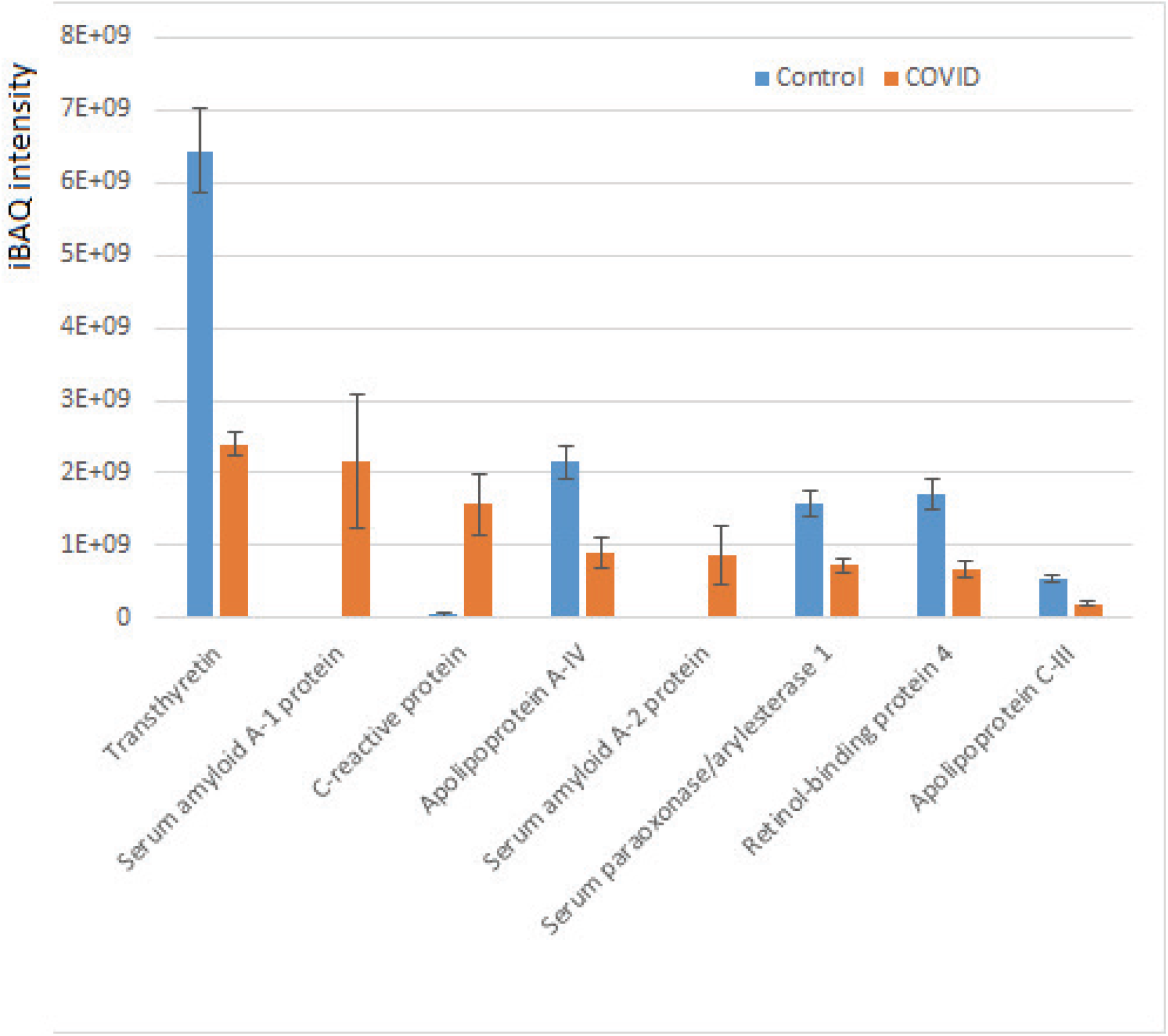
Plasma proteins from COVID-19 patients exhibits prominent changes compared to control. iBAQ intensities of the most abundant proteins showing significant differences between COVID-19 and control. See Fig. S4 for the full list of hits. Bars represent normalized intensity levels averaged for all samples in each group, error bars represent SEM (n=7 for control, 6 for COVID-19).

Thus, our data suggest that a limited set of proteins related to inflammation, immune response, and normal organismal homeostasis are prominently altered between COVID-19 and control, potentially as a direct consequence of SARS-CoV-2 infection.

### COVID-19 plasma proteome exhibits altered posttranslational modification patterns

To test whether COVID-19 is associated with any changes in PTMs, we analyzed our total proteomics run of plasma samples using pFIND, a software package that can simultaneously identify a large number of different PTMs and chemical modifications based on mass shifts ^21,22^. We manually added arginylation into the program (including addition of unmodified, mono- and dimethyl-Arg that have been all shown to occur *in vivo* through our previous work ^23-25^). This search identified a total of 2630 modifications in the samples. The results were filtered by precursor mass tolerance of 10 ppm, fragment ion tolerance of 20 ppm, and false discovery rate (FDR) <1% at peptide and protein level.

Total results from the pFIND search, including data in individual samples, is shown in Table S5, and the 10 most abundant modifications from COVID-19 and control patient samples are plotted in Fig. S4. These results were used to calculate each modified peptide amount as spectra count normalized to the percent of the total peptides in the sample, and these numbers were then compared across samples between control and COVID-19 to calculate p-value and fold change.

We defined putative significant differences as a greater than 1.5 fold change between control and COVID-19 peptides, with p-value less than 0.05. These hits are listed in Table S6, and the full list of modifications and modified sites identified in our search is listed in Table S7.

A total of 82 PTMs showed statistically significant differences between COVID-19 and control. However, many of these modifications are apparently chemically induced and have not been described to happen under normal physiological conditions. Such modifications in the plasma can potentially occur in response to drugs and environmental factors; thus, it is unlikely that these modifications occur as a result of SARS-CoV-2 infection, even though they might be directly or indirectly related to the patients’ treatment or susceptibility to symptomatic COVID-19. Given this uncertainty, we excluded these PTMs from further analysis, and manually selected only the known naturally occurring modifications for further analysis.

After this filtering, only a few physiological PTMs showed significant differences f between COVID-19 and controls (Fig. 4). These PTMs were plotted in three separate groups: high abundance (Fig. 4A), intermediate abundance (Fig. 4B), and low abundance (Fig. 4C).Notably, only three of them were in the high abundance group, including phosphorylation on Thr, which showed a nearly 2-fold increase in COVID-19 patients, Arg deamidation, ∼2-fold increased in control, and a >1.5-fold reduction in side chain arginylation of Asp and Glu residues in COVID-19 patients (Fig. 4A). Arginylation is an emerging poorly characterized modification, which is still non-routine during proteomics analysis and normally requires manual data validation. While it was not feasible to manually validate all identified peptides, we validated a number of representative MS/MS spectra (Dataset 1) to confirm that arginylation on the identified sites is likely.

**Figure 4.**
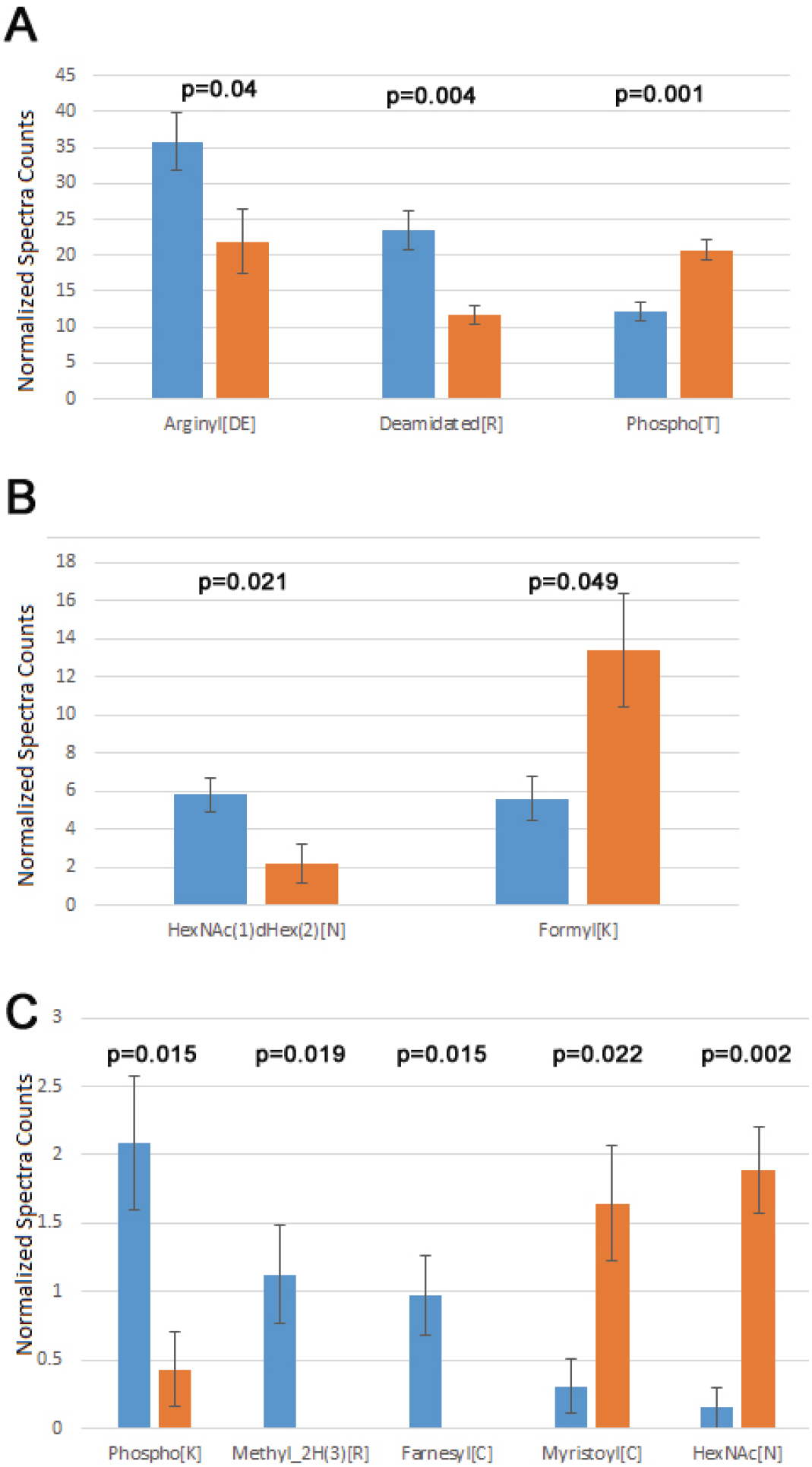
Plasma proteins from COVID-19 patients exhibits changes in the overall levels of several physiological posttranslational modifications. Normalized spectra counts for high (A), intermediate (B), and low abundance (C) hits are plotted on different scales. Error bars represent SEM (n=7 for control, 6 for COVID-19). P values calculated by 2-tailed Student’s T-test are listed on top of each set of bars.

Both arginylation and Thr phosphorylation have been previously proposed to play a global regulatory role. Thus, we performed further analysis of these two modifications to dissect the likely functional consequences of their change in COVID-19. Mapping the identified sites for these two modifications on the target proteins identified in our sets (Tables 1 and 2) revealed that in addition of the total level change for each of the PTMs, COVID-19 patients also exhibit a different repertoire of arginylation and phosphorylation sites compared to control. This points to a likely possibility that the target proteins affected by these altered regulatory PTMs, including blood coagulation and immune response, along with several other functions implicated in COVID-19 infection, may exhibit different behavior in response to SARS-CoV-2. It is attractive to suggest that arginylation/phosphorylation on these sites may underlie COVID-19 dependent protein regulation and disease progression.

The two less abundant lists included a number of additional PTMs that play important physiological roles. Among these were three types of glycosylation (N-linked up in controls, and S-linked and N-terminal up in COVID-19). Notably, changes in glycosylation were previously observed on the viral SARS-CoV-2 proteins during COVID-19^26-29^, even though such changes in the host organism, to the best of our knowledge, were not previously reported. Additional modifications on this list were detected at very low levels. Even though physiologically important, it is difficult to assess the biological role of their changes in the plasma of COVID-19 patients.

In addition to these PTMs, a substantial number of amino acid substitutions were significantly different between COVID-19 and control plasma (Fig S5). Of those, Glu to Asp substitution was by far the most abundant (Fig. S6, bottom). These substitutions normally reflect single nucleotide polymorphisms (SNPs), and thus are not a result of SARS-CoV-2 infection. However, the presence of these SNPs potentially reflect genetic changes that might make the patients more vulnerable to COVID-19.

## Conclusions

To our knowledge, the data presented in this manuscript represents the first global proteome and peptidome analysis of plasma from COVID-19 patients. Other studies have extensively analyzed the proteomics and posttranslational state of SARS-CoV-2 proteins during COVID-19 infection, but no one has as yet focused on the changes in the patient plasma proteome.

Our data suggests the existence of global protein-level trends that may inform our understanding of biology, prevention, diagnostics, and treatment of COVID-19. We find that plasma proteins in COVID-19 patients contain elevated levels of a limited number of proteins associated with inflammation and immune response, and some of these proteins, including fibrinogen, apparently undergo different proteolytic events compared to normal controls. Some of these changes may underlie the less understood clinical symptoms of COVID-19.

Our study shows that COVID-19 infection is accompanied by a prominent posttranslational signature, with differential increase and reduction in several key regulatory modifications, including glycosylation, citrullination, Thr phosphorylation, and arginylation. Decoding this signature may potentially inform our understanding of the biology, diagnostics, and treatment of COVID-19.

## Materials and methods

### Human patient samples

Blood from 7 healthy controls and 7 COVID-19 patients was collected in lavender-top EDTA tubes from BD (cat no. 368661). These tubes come with 10.8 mg K2 EDTA spray dried inside the tube. Blood was drawn into the tube by clinical staff and handed off to the processing unit within 8 hours, usually much less. Tubes were then spun down at 1000 x g at room temperature for 15 minutes. Plasma supernatant was removed, aliquoted, snap frozen and stored on dry ice for ∼1 hour, then transferred to -80°C for storage.

### Plasma fractionation for protein and peptide fractions

#### Plasma clarification

Frozen plasma aliquots were thawed on ice and centrifuged at 13,000g at 4°C for 30 minutes. The supernatant was used for peptide isolation or further depleted of IgG and albumin for analyzing the protein composition using mass spectrometry.

#### Peptide isolation

50µl of clarified plasma was mixed with 200 µl of Phosphate Buffered Saline. 1 ml of freshly prepared 100 mM Tris, 8 M urea, pH 7.5 at room temperature was used to denature the proteins and dissociate bound peptides from abundant plasma proteins. Denatured samples were loaded onto a pre-rinsed Amicon Ultra 30kDa MWCO filter (Millipore) and centrifuged at 13,000g at 4°C for 20 minutes. Formic acid was added to the flow-through containing primarily peptides to a final concentration of 0.1% (v/v). MacroSpin Vydac Silica C18 columns (the Nest Group, Inc, Part #SMM SS18V, Lot #060310) were pre-washed first with 500µl of 100% acetonitrile and then with water followed by centrifugation at 100g at 4°C for 1 minute to remove the liquid after each wash. The same centrifugation condition was used for all subsequent steps. Briefly, columns were equilibrated with 300µl 0.2% acetonitrile; the 30 kDA MWCO flow-through was applied; columns were washed three times with 400µl of 0.1% formic acid, and peptides were then eluted using 300µl of 0.1% formic acid, 50% acetonitrile, snap frozen in liquid nitrogen, and stored at -80°C until the analysis.

#### Albumin & IgG depletion for proteomics

The plasma samples were depleted of IgG and albumin using Albumin & IgG depletion SpinTrap columns (GE healthcare) according to manufacturer’s instructions. Briefly, the column was resuspended, and the storage buffer was discarded by centrifugation at 100g for 30 seconds. The column was then equilibrated with 400µl of binding buffer (20mM sodium phosphate, 150mM sodium chloride, pH 7.4), which was discarded by centrifugation at 800g for 30 seconds. Immediately after thawing an aliquot, the plasma sample was diluted 1:1 with binding buffer, applied to the column, mixed with the resin and incubated for 5 minutes at room temperature. The albumin and IgG depleted protein fraction was collected by centrifugation at 800g for 30 seconds, the column was washed twice with 100µl binding buffer flowed by centrifugation at 800g for 30 seconds, and the flowthrough and washes were combined. The eluted proteins were precipitated by adding 9 volumes of -20°C ethanol and stored overnight at 4°C. Precipitated protein was collected by centrifugation at 13000g at 4°C for 30 minutes. The pellet was heat inactivated at 60°C for 20 minutes to denature any viral load that might have been present.

### Peptidomics

#### Sample Preparation and LC-MS/MS analysis

Total peptide fraction purified by C18 column from 13 COVID-19 patient and control plasma (7 controls and 6 diseased) were lyophilized and resuspended in 30 ul of 3% acetonitrile, 0.1% formic acid. The volume of plasma was kept constant between samples at each step to enable non-normalized comparison of abundances for each identified peptide. 5 μl of each sample was analyzed by LC-MS/MS on the Thermo Q Exactive HF mass spectrometer using a 2-h LC gradient^30^.

#### Data Analysis

MS/MS data were analyzed using Thermo Proteome Discoverer v2.4. Spectra were searched using no-enzyme specificity against the UniProt human proteome database (10/02/2020) and a common contaminant database using Sequest HT. Percolator target False Discovery Rate was set at 0.01 (Strict) and 0.05 (Relaxed). Only peptides identified with high confidence were retained. Common contaminants (primarily keratins) were removed. Peptide abundance values were determined from the peptide chromatographic peak areas. Ratio, p-value (t-test) and adjusted q-value (p-value adjusted to account for multiple testing using Benjamini-Hochberg FDR) were calculated using the non-normalized abundance values. Significant changing peptides are defined as peptides with a minimum absolute fold change of 2, adjusted p-value of 0.05, and identified in a minimum of 3 replicates in either group.

To produce the charts shown in the main and supplemental figures, we summed up total abundances of all peptides belonging to each protein that exhibited significant differences between groups and calculated the average of these sums between all control and all COVID-19 samples. For the chart showing differential proteolytic patterns of fibrinogen and serglycin (Fig. 2), abundances of individual peptides were averaged across samples and plotted against each peptide sequence.

### Proteomics

#### Sample Preparation and LC-MS/MS analysis

The ethanol precipitated albumin and IgG depleted COVID-19 patient plasma (7 controls, denoted by 3-digit numbers starting with 0 in the supplemental tables, and 6 diseased, denoted by 3-digit numbers starting with 5 in the supplemental tables) fractions were dissolved in 50 μl of 1% SDS, 50 mM Tris-Cl pH 7.5. 10 μl of each was run into a NuPAGE 10% Bis-Tris gel (Thermo Scientific) for a short distance. The entire stained gel regions were excised, reduced with tris(2-carboxyethyl)phosphine (TCEP), alkylated with iodoacetamide, and digested with trypsin. Tryptic digests were analyzed using a single-shot extended 4h LC gradient on the Thermo Q Exactive Plus mass spectrometer.

#### Data Analysis

Peptide sequences were identified using MaxQuant 1.6.17.0 (Ref: PMID 19029910). MS/MS spectra were searched against a UniProt human proteome database (10/02/2020) and a common contaminants database using full tryptic specificity with up to two missed cleavages, static carboxamidomethylation of Cys, and variable Met oxidation, protein N-terminal acetylation and Asn deamidation. “Match between runs” feature was used to help transfer identifications across experiments to minimize missing values. Consensus identification lists were generated with false discovery rates set at 1% for protein and peptide identifications. Statistical analyses were performed using Perseus 1.6.15.0 (Ref: PMID 27348712). Protein fold changes were determined from the Intensity values. Missing values were imputed with a minimum Intensity value, and t-test p-values were adjusted to account for multiple testing using the permutation-based FDR function in Perseus. High confidence identification of proteins with significant change was determined based on the following criteria: minimum absolute fold change of 2, q-value <0.05, identified by a minimum of 2 razor + unique peptides, and detected in at least 3 of the replicates in one of the groups compared.

### Posttranslational modifications analysis

The “Proteomics” MS/MS data described above were analyzed using pFind 3.1.5^31^. Two separate pFind searches were performed: Control (containing all 7 control samples) and Disease (containing all 6 samples). Spectra were searched using partial tryptic specificity against the UniProt human proteome database (10/02/2020). The “Open Search” option was used to identify PTMs. Data were filtered using a precursor mass tolerance of 10 ppm, fragment ion tolerance of 20 ppm, and FDR <1% at peptide and protein level.

### Data deposition

All mass spectrometry raw data have been uploaded to the MassIVE public repository (https://massive.ucsd.edu/ProteoSAFe/static/massive.jsp) with the accession MSV000088382, and the ProteomeXchange repository (http://www.proteomecentral.proteomexchange.org/cgi/) with the accession PXD029756.

## Acknowledgements

We are grateful to Dr. John Wherry and Penn Immune Health Initiative for support of this work and providing the plasma samples for this analysis. We thank the Human Immunology Core (HIC) facility for assistance with sample processing and provision of control samples for this study. The HIC is funded in part by NIH P30-AI0450080 and P30-CA016520. This work was supported by the NIH grant R35GM122505 to AK and R50 CA221838 to HYT.

## Author contributions

PV, AK, HYT, and DS designed the research

PV and HYT performed the experiments and analyzed data

AK and DS analyzed data and wrote the paper

CPU collected and provided materials for analysis

## References

1. Swelum AA, Shafi ME, Albaqami NM, et al. COVID-19 in Human, Animal, and Environment: A Review. Front Vet Sci. 2020;7:578. doi: 10.3389/fvets.2020.00578.

2. Novelli G, Biancolella M, Mehrian-Shai R, et al. COVID-19 one year into the pandemic: from genetics and genomics to therapy, vaccination, and policy. Hum Genomics. 2021;15(1):27. doi: 10.1186/s40246-021-00326-3.

3. Ostergaard L. SARS CoV-2 related microvascular damage and symptoms during and after COVID-19: Consequences of capillary transit-time changes, tissue hypoxia and inflammation. Physiol Rep. 2021;9(3):e14726. doi: 10.14814/phy2.14726.

4. Harapan BN, Yoo HJ. Neurological symptoms, manifestations, and complications associated with severe acute respiratory syndrome coronavirus 2 (SARS-CoV-2) and coronavirus disease 19 (COVID-19). J Neurol. 2021;268(9):3059–3071. doi: 10.1007/s00415-021-10406-y.

5. Vakil-Gilani K, O’Rourke K. Are patients with rheumatologic diseases on chronic immunosuppressive therapy at lower risk of developing severe symptoms when infected with COVID-19? Clin Rheumatol. 2020;39(7):2067–2068. doi: 10.1007/s10067-020-05184-3.

6. Schmulson M, Davalos MF, Berumen J. Beware: Gastrointestinal symptoms can be a manifestation of COVID-19. Rev Gastroenterol Mex (Engl Ed). 2020;85(3):282–287. doi: 10.1016/j.rgmx.2020.04.001.

7. Troyer EA, Kohn JN, Hong S. Are we facing a crashing wave of neuropsychiatric sequelae of COVID-19? Neuropsychiatric symptoms and potential immunologic mechanisms. Brain Behav Immun. 2020;87:34–39. doi: 10.1016/j.bbi.2020.04.027.

8. Hanff TC, Mohareb AM, Giri J, Cohen JB, Chirinos JA. Thrombosis in COVID-19. Am J Hematol. 2020;95(12):1578–1589. doi: 10.1002/ajh.25982.

9. Arapidi G, Osetrova M, Ivanova O, et al. Peptidomics dataset: Blood plasma and serum samples of healthy donors fractionated on a set of chromatography sorbents. Data Brief. 2018;18:1204–1211. doi: 10.1016/j.dib.2018.04.018.

10. Meyer B, Chiaravalli J, Gellenoncourt S, et al. Characterising proteolysis during SARS-CoV-2 infection identifies viral cleavage sites and cellular targets with therapeutic potential. Nat Commun. 2021;12(1):5553. doi: 10.1038/s41467-021-25796-w.

11. Anand P, Puranik A, Aravamudan M, Venkatakrishnan AJ, Soundararajan V. SARS-CoV-2 strategically mimics proteolytic activation of human ENaC. Elife. 2020;9. doi: 10.7554/eLife.58603.

12. Ramos-Guzman CA, Ruiz-Pernia JJ, Tunon I. Unraveling the SARS-CoV-2 Main Protease Mechanism Using Multiscale Methods. ACS Catal. 2020;10:12544–12554. doi: 10.1021/acscatal.0c03420.

13. Swiderek K, Moliner V. Revealing the molecular mechanisms of proteolysis of SARS-CoV-2 M(pro) by QM/MM computational methods. Chem Sci. 2020;11(39):10626–10630. doi: 10.1039/d0sc02823a.

14. Tang T, Jaimes JA, Bidon MK, Straus MR, Daniel S, Whittaker GR. Proteolytic Activation of SARS-CoV-2 Spike at the S1/S2 Boundary: Potential Role of Proteases beyond Furin. ACS Infect Dis. 2021;7(2):264–272. doi: 10.1021/acsinfecdis.0c00701.

15. Katneni UK, Alexaki A, Hunt RC, et al. Coagulopathy and Thrombosis as a Result of Severe COVID-19 Infection: A Microvascular Focus. Thromb Haemost. 2020;120(12):1668–1679. doi: 10.1055/s-0040-1715841.

16. Gomez-Mesa JE, Galindo-Coral S, Montes MC, Munoz Martin AJ. Thrombosis and Coagulopathy in COVID-19. Curr Probl Cardiol. 2021;46(3):100742. doi: 10.1016/j.cpcardiol.2020.100742.

17. Montenegro F, Unigarro L, Paredes G, et al. Acute respiratory distress syndrome (ARDS) caused by the novel coronavirus disease (COVID-19): a practical comprehensive literature review. Expert Rev Respir Med. 2021;15(2):183–195. doi: 10.1080/17476348.2020.1820329.

18. May JE, Wolberg AS, Lim MY. Disorders of Fibrinogen and Fibrinolysis. Hematol Oncol Clin North Am. 2021. doi: 10.1016/j.hoc.2021.07.011.

19. Ahmed S, Zimba O, Gasparyan AY. Thrombosis in Coronavirus disease 2019 (COVID-19) through the prism of Virchow’s triad. Clin Rheumatol. 2020;39(9):2529–2543. doi: 10.1007/s10067-020-05275-1.

20. Scully OJ, Chua PJ, Harve KS, Bay BH, Yip GW. Serglycin in health and diseases. Anat Rec (Hoboken). 2012;295(9):1415–1420. doi: 10.1002/ar.22536.

21. Wang LH, Li DQ, Fu Y, et al. pFind 2.0: a software package for peptide and protein identification via tandem mass spectrometry. Rapid Commun Mass Spectrom. 2007;21(18):2985–2991. doi: 10.1002/rcm.3173.

22. Li D, Fu Y, Sun R, et al. pFind: a novel database-searching software system for automated peptide and protein identification via tandem mass spectrometry. Bioinformatics. 2005;21(13):3049–3050. doi: 10.1093/bioinformatics/bti439.

23. Wang J, Yates JR 3rd,, Kashina A. Biochemical analysis of protein arginylation. Methods Enzymol. 2019;626:89–113. doi: 10.1016/bs.mie.2019.07.028.

24. Wang J, Han X, Wong CC, et al. Arginyltransferase ATE1 catalyzes midchain arginylation of proteins at side chain carboxylates in vivo. Chem Biol. 2014;21(3):331–337. doi: 10.1016/j.chembiol.2013.12.017.

25. Saha S, Wong CC, Xu T, et al. Arginylation and methylation double up to regulate nuclear proteins and nuclear architecture in vivo. Chem Biol. 2011;18(11):1369–1378. doi: 10.1016/j.chembiol.2011.08.019.

26. Zhong L, Zhu L, Cai ZW. Mass Spectrometry-based Proteomics and Glycoproteomics in COVID-19 Biomarkers Identification: A Mini-review. J Anal Test. 2021:1–16. doi: 10.1007/s41664-021-00197-6.

27. Shajahan A, Pepi LE, Rouhani DS, Heiss C, Azadi P. Glycosylation of SARS-CoV-2: structural and functional insights. Anal Bioanal Chem. 2021. doi: 10.1007/s00216-021-03499-x.

28. Praissman JL, Wells L. Proteomics-Based Insights Into the SARS-CoV-2-Mediated COVID-19 Pandemic: A Review of the First Year of Research. Mol Cell Proteomics. 2021;20:100103. doi: 10.1016/j.mcpro.2021.100103.

29. Reis CA, Tauber R, Blanchard V. Glycosylation is a key in SARS-CoV-2 infection. J Mol Med (Berl). 2021;99(8):1023–1031. doi: 10.1007/s00109-021-02092-0.

30. Su C, Lu F, Soldan SS, et al. EBNA2 driven enhancer switching at the CIITA-DEXI locus suppresses HLA class II gene expression during EBV infection of B-lymphocytes. PLoS Pathog. 2021;17(8):e1009834. doi: 10.1371/journal.ppat.1009834.

31. Chi H, Liu C, Yang H, et al. Comprehensive identification of peptides in tandem mass spectra using an efficient open search engine. Nat Biotechnol. 2018. doi: 10.1038/nbt.4236.

