## Supplementary material for "Protein Posttranslational Signatures Identified in COVID-19 Patient Plasma": Dataset 1

Selected LC-MS/MS spectra of arginylated peptides found in COVID-19 and control plasma samples

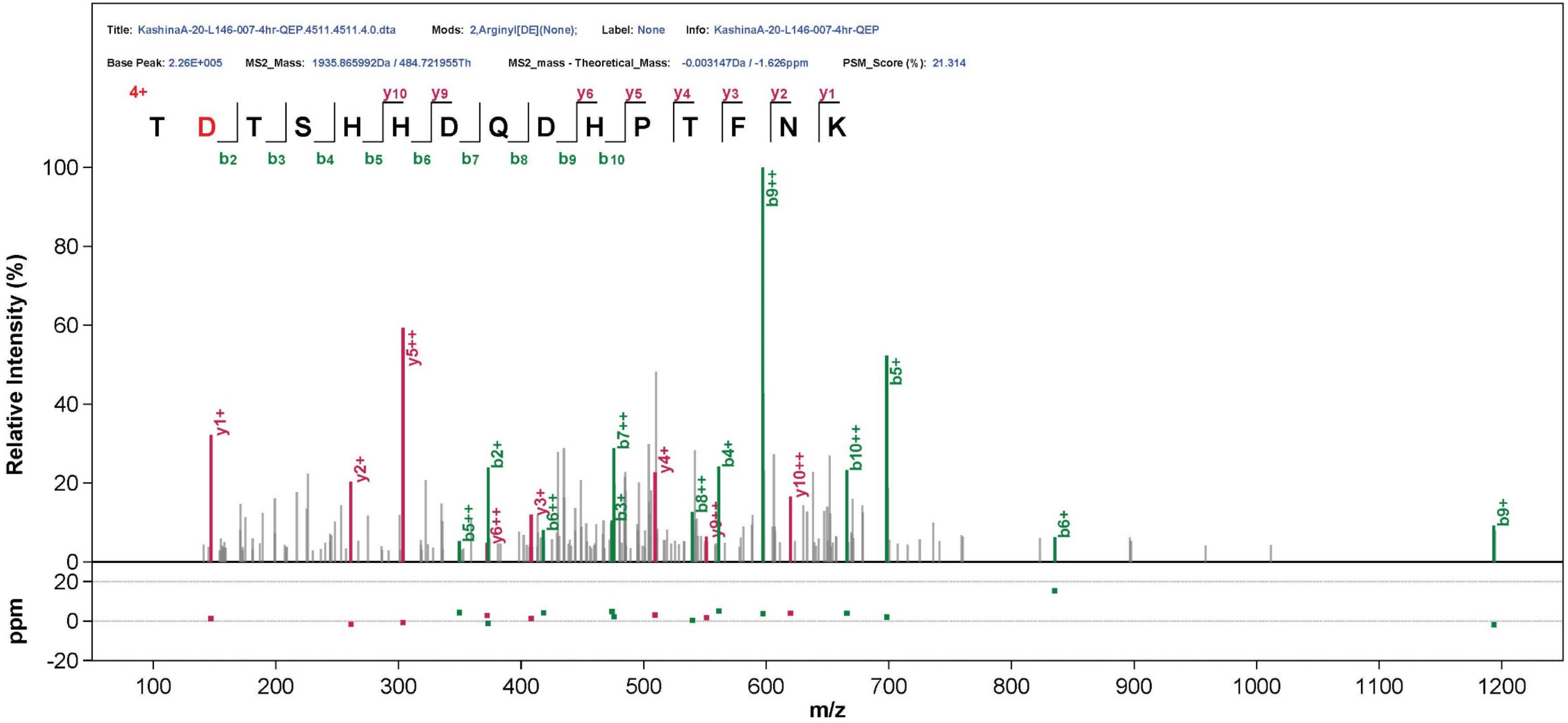

Title: KashinaA-20-L146-007-4hr-QEP.4511.4511.4.0.dta
Mods: 2,Arginyl[DE](None);
Label: None
Info: KashinaA-20-L146-007-4hr-QEP
Base Peak: 2.26E+005
MS2_Mass: 1935.865992Da / 484.721955Th
MS2_mass - Theoretical_Mass: -0.003147Da / -1.626ppm
PSM_Score (%): 21.314
4+
T D T S H H D Q D H P T F N K
Relative Intensity (%)
ppm
m/z

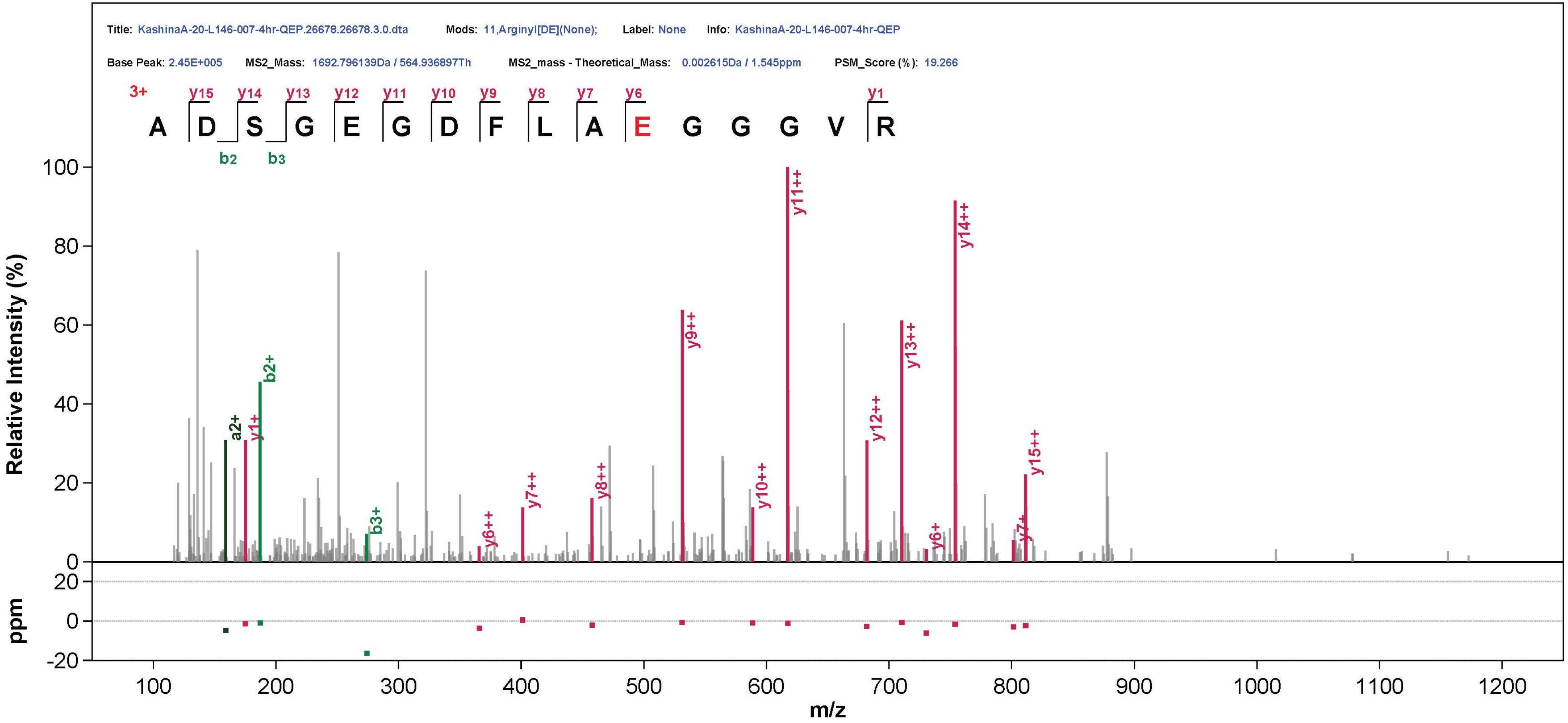

Title: KashinaA-20-L146-007-4hr-QEP.26678.26678.3.0.dta
Mods: 11,Arginyl[DE](None);
Label: None
Info: KashinaA-20-L146-007-4hr-QEP
Base Peak: 2.45E+005
MS2_Mass: 1692.796139Da / 564.936897Th
MS2_mass - Theoretical_Mass: 0.002615Da / 1.545ppm
PSM_Score (%): 19.266
3+
A D S G E G D F L A E G G G V R
Relative Intensity (%)
ppm
m/z

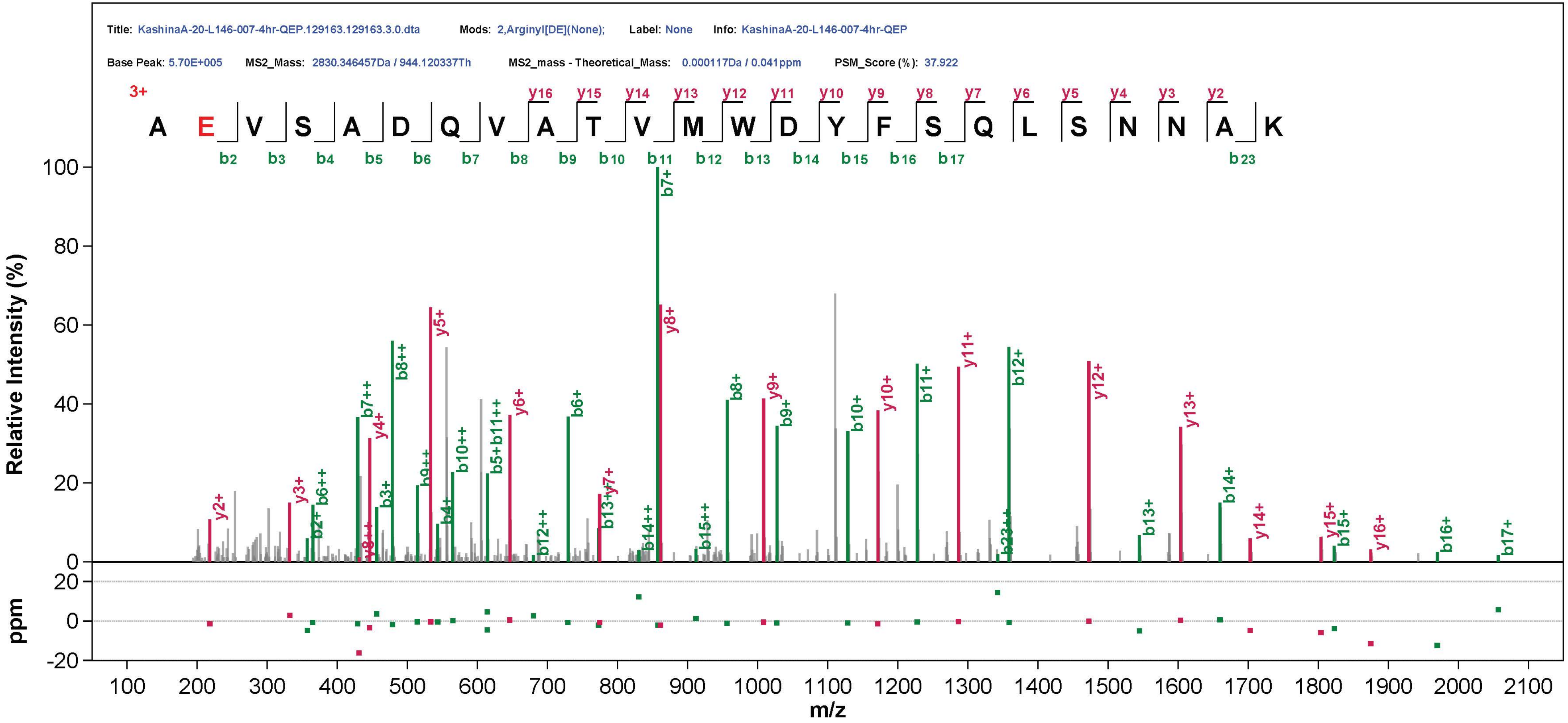
Title: KashinaA-20-L146-007-4hr-QEP.129163.129163.3.0.dta
Mods: 2,Arginyl[DE](None);
Label: None
Info: KashinaA-20-L146-007-4hr-QEP
Base Peak: 5.70E+005
MS2_Mass: 2830.346457Da / 944.120337Th
MS2_mass - Theoretical_Mass: 0.000117Da / 0.041ppm
PSM_Score (%): 37.922
3+
A E V S A D Q V A T V M W D Y F S Q L S N N A K
Relative Intensity (%)
ppm
m/z

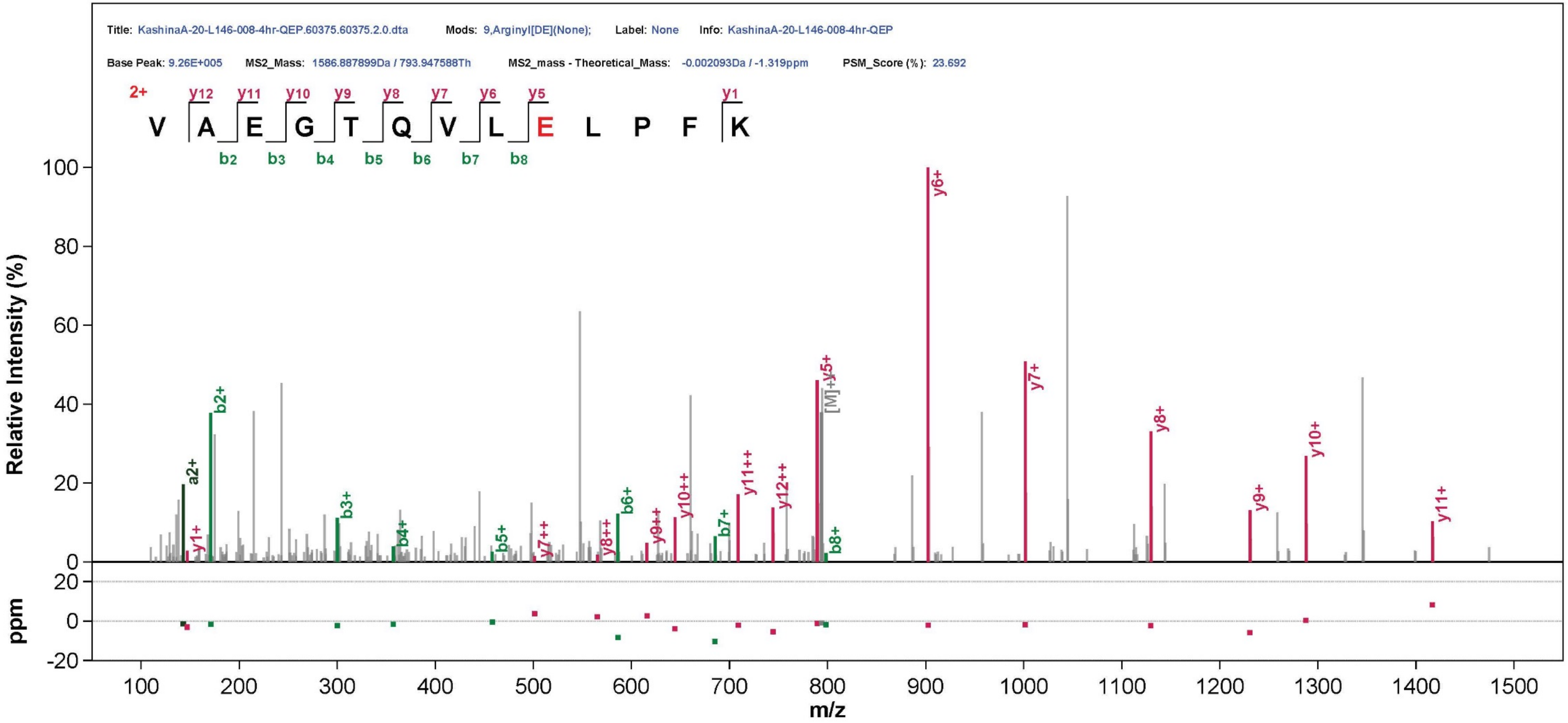

Title: KashinaA-20-L146-008-4hr-QEP.60375.60375.2.0.dta
Mods: 9,Arginyl[DE](None);
Label: None
Info: KashinaA-20-L146-008-4hr-QEP
Base Peak: 9.26E+005
MS2_Mass: 1586.887899Da / 793.947588Th
MS2_mass - Theoretical_Mass: -0.002093Da / -1.319ppm
PSM_Score (%): 23.692
2+
V A E G T Q V L E L P F K
Relative Intensity (%)
ppm
m/z

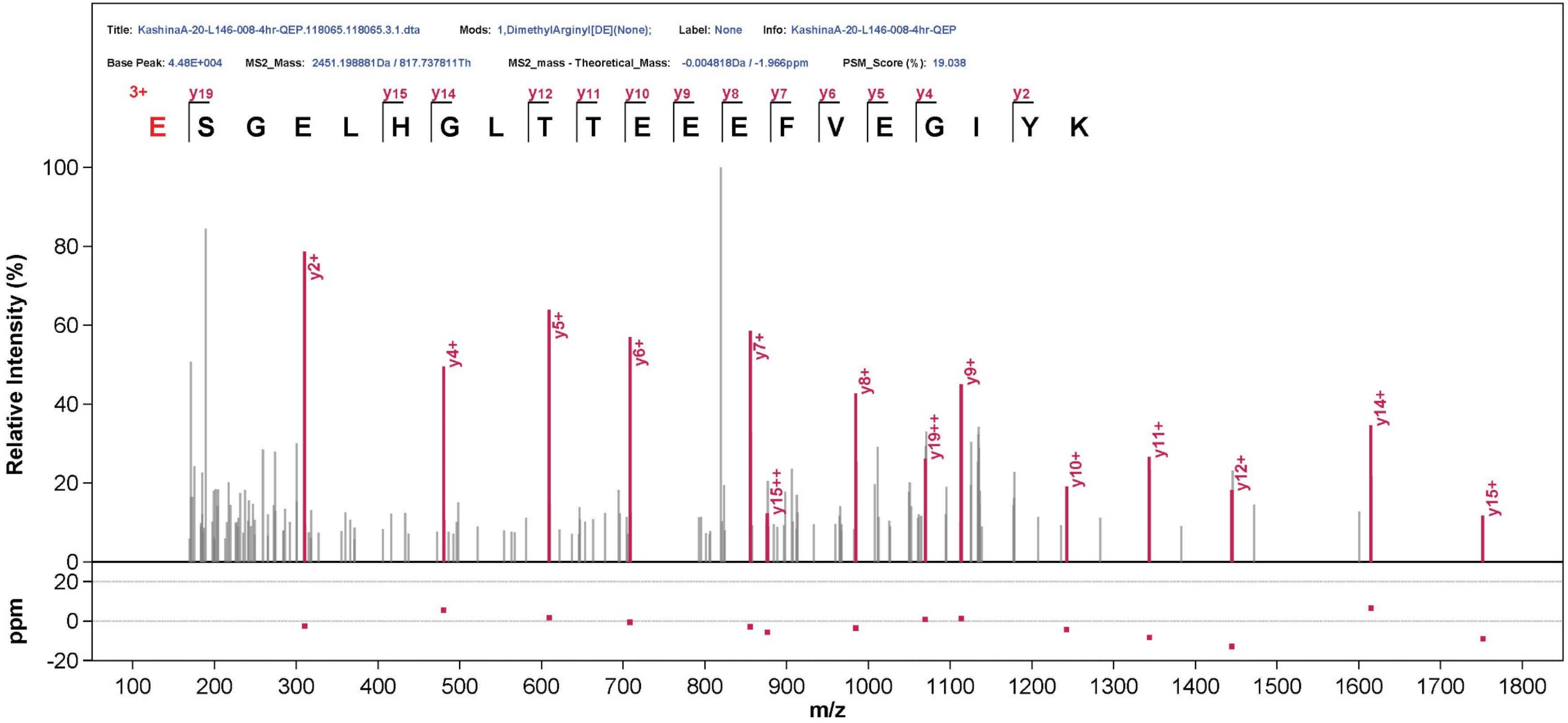
Title: KashinaA-20-L146-008-4hr-QEP.118065.118065.3.1.dta
Mods: 1,DimethylArginyl[DE](None);
Label: None
Info: KashinaA-20-L146-008-4hr-QEP
Base Peak: 4.48E+004
MS2_Mass: 2451.198881Da / 817.737811Th
MS2_mass - Theoretical_Mass: -0.004818Da / -1.966ppm
PSM_Score (%): 19.038
3+
E S G E L H G L T T E E E F V E G I Y K
y19 y15 y14 y12 y11 y10 y9 y8 y7 y6 y5 y4 y2
Relative Intensity (%)
y2+
y4+
y5+
y6+
y7+
y15++
y8+
y19++
y9+
y10+
y11+
y12+
y14+
y15+
ppm
m/z

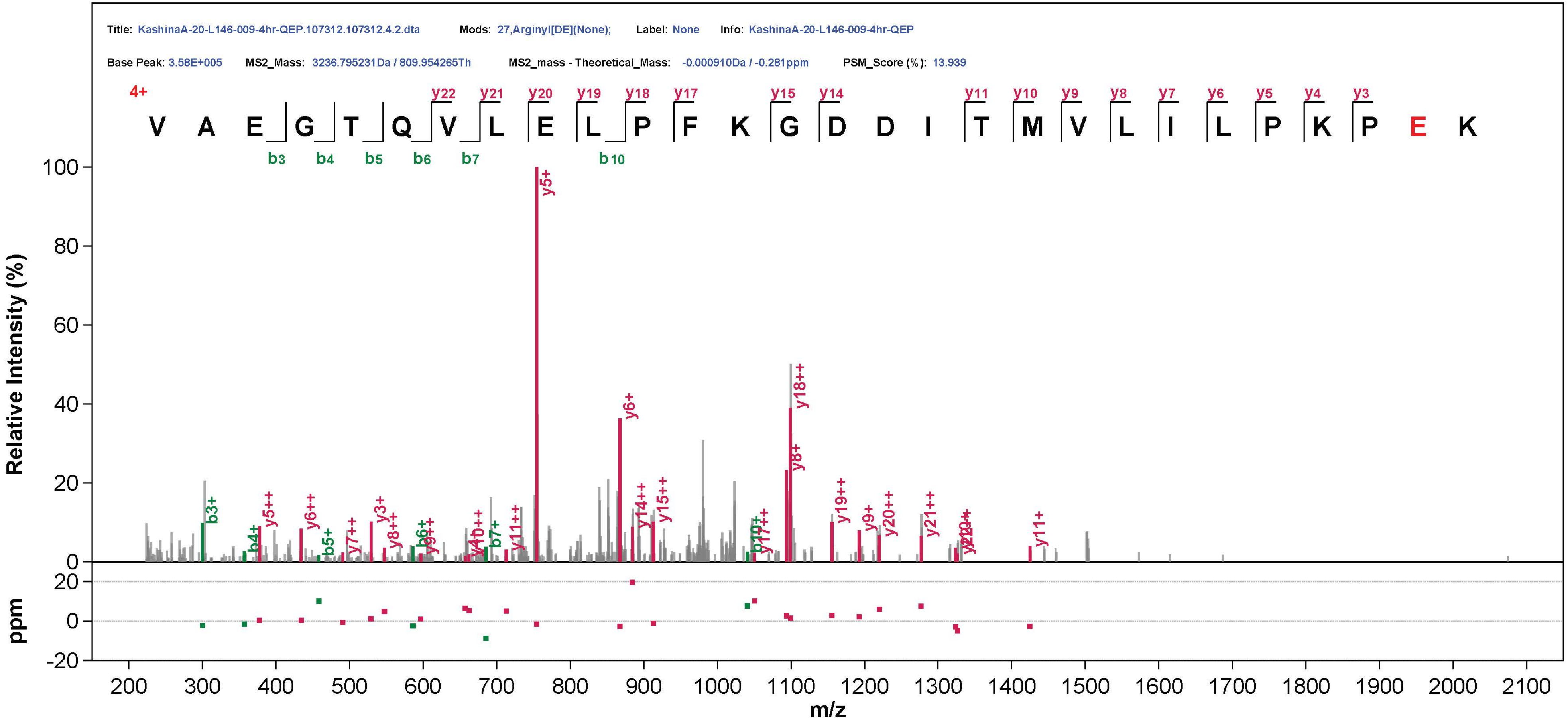

Title: KashinaA-20-L146-009-4hr-QEP.107312.107312.4.2.dta
Mods: 27,ArginylI[DE](None);
Label: None
Info: KashinaA-20-L146-009-4hr-QEP
Base Peak: 3.58E+005
MS2_Mass: 3236.795231Da / 809.954265Th
MS2_mass - Theoretical_Mass: -0.000910Da / -0.281ppm
PSM_Score (%): 13.939
4+
V A E G T Q V L E L P F K G D D I T M V L I L P K P E K
Relative Intensity (%)
ppm
m/z
