## Supplementary material for "Protein Posttranslational Signatures Identified in COVID-19 Patient Plasma": Table S1

| **Sample ID** | **Gender** | **Age** | **Collection Date** | **COVID-19 diagnosis** | **Clots** |
| --- | --- | --- | --- | --- | --- |
| 007 | F | 40 | 4/13/20 | Negative | No |
| 008 | F | 40 | 4/20/20 | Negative | No |
| 009 | M | 44 | 4/20/20 | Negative | No |
| 010 | F | 36 | 4/22/20 | Negative | No |
| 014 | F | 57 | 4/28/20 | Negative | No |
| 017 | M | 41 | 4/30/20 | Negative | No |
| 027 | F | 49 | 4/20/05 | Negative | No |
| 548 | M | 39 | 4/17/20 | Positive | Yes |
| 559 | F | 54 | 4/21/20 | Positive | Yes |
| 571 | F | 49 | 4/28/20 | Positive | Yes |
| 562 | F | 30 | 4/23/20 | Positive | No |
| 540 | F | 44 | 4/15/20 | Positive | No |
| 558 | F | 48 | 4/21/20 | Positive | No |
