## Supplementary material for "Protein Posttranslational Signatures Identified in COVID-19 Patient Plasma": Table S8

| **Name** | **Gene symbol** | **Uniprot Accession** | **Modification Site Specific to Control** | **Modification Site Specific to COVID** | **Modification Site Common** | **Function** |
| --- | --- | --- | --- | --- | --- | --- |
| Afamin | AFAM | P43652 | E186; E369 |  | D364 | Carrier for hydrophobic molecules in body fluids |
| Alpha-1-acid glycoprotein 2 | A1AG2 | P19652 |  | D172 |  | Transport protein for hydrophobic ligands in the blood stream |
| Alpha-1-antichymotrypsin | AACT | P01011 |  |  | E285; E286 | Protease inhibitor |
| Alpha-1-antitrypsin | A1AT | P01009 | D201 | E219; D322 | D36; E146: E348 | Protease inhibitor |
| Alpha-2-macroglobulin | A2MG | P01023 | D134; E216; E624; D672; E1198 |  |  | Protease inhibitor |
| Antithrombin-III | ANT3 | P01008 | D65 |  | E303; E321 | Blood coagulation and clot formation |
| Apolipoprotein A4 | APOA4 | P06727 | E21; E312; E323 |  | D213; E291 | Fat metabolism and lipid transport |
| Apolipoprotein A-I | APOA1 | P02647 | E100; E229 |  | D25; E26; D33; D44; D192; E193 | Cholesterol efflux |
| Apolipoprotein B-100 | APOB | P04114 |  |  | E172 | Fat metabolism and lipid transport |
| Apolipoprotein E | APOE | P02649 | E31 |  |  | Fat metabolism; lipid transport between organs; link to neurodegeneration; gender-specific |
| Ceruloplasmin | CERU | P00450 | E731; E594 | E615 | D933 | Iron transport across the cell membrane |
| Complement C3 | CO3 | P01024 | E294; D1417 |  |  | Complement pathway, immune response |
| Complement C4-B | CO4B | P0C0L5 |  |  | D146; D860; D1098 | Complement pathway, immune response |
| Complement factor H | CFAH | P08603 |  |  | E44 | Complement pathway, immune response |
| Fibrinogen alpha chain | FIBA | P02671 | E30; D199; E389; D396; D515; E559; E606 | D174; E545; E610 | E284 | Blood coagulation and clot formation |
| Fibrinogen beta chain | FIBB | P02675 | D419 | E177; D327 | E185; D346 | Blood coagulation and clot formation |
| Fibrinogen gamma chain | FIBG | P02679 | D130; D229 |  | E257 | Blood coagulation and clot formation |
| Ficolin-3 | FCN3 | O75636 |  |  | D142 | Complement pathway, immune response |
| Haptoglobin | HPT | P00738 |  |  |  | Combines with free plasma hemoglobin to allow hepatic recycling of heme iron and to prevent kidney damage |
| Hemoglobin subunit alpha | HBA | P69905 |  | D86 |  | Oxygen transport |
| Hemoglobin subunit beta | HBB | P68871 |  | D80 |  | Oxygen transport |
| Immunoglobulin heavy constant mu | IGHM | P01871 | E167; D330 |  |  | Antibody |
| Immunoglobulin heavy variable 3-11 | HV311 | P01762 | D81 |  |  | Antibody |
| Immunoglobulin kappa constant | IGKC | P01834 |  |  | D44; D60 | Antibody |
| Immunoglobulin lambda variable 3-21 | LV321 | P80748 |  |  | D69 | Antibody |
| Inter-alpha-trypsin inhibitor heavy chain H4 | ITIH4 | Q14624 | E143; E332 |  | E148 | Protease inhibitor |
| N-acetylmuramoyl-L-alanine amidase | PGRP2 | Q96PD5 |  |  | D164 | scavenger role by digesting biologically active peptidoglycan (PGN) into biologically inactive fragments |
| Serotransferrin | TRFE | P02787 |  | D216; E300 | D461 | Iron transport from sites of absorption and heme degradation to those of storage and utilization |
| Serum amyloid A-2 protein | SAA2 | P0DJI9 |  | E74 |  | Apolipoproteins associated with high-density lipoprotein in plasma |
| Transthyretin | TTHY | P02766 | E71 |  |  | Thyroid hormone-binding protein |
| von Willebrand factor | VWF | P04275 | D1622 |  |  | Blood coagulation and clot formation |
