## Supplementary material for "Protein Posttranslational Signatures Identified in COVID-19 Patient Plasma": Table S9

| **Name** | **Gene symbol** | **Uniprot Accession** | **Modification Site Specific to Control** | **Modification Site Specific to COVID** | **Modification Site Common** | **Function** |
| --- | --- | --- | --- | --- | --- | --- |
| Alpha-1-antichymotrypsin | AACT | P01011 |  |  | T35 | Protease inhibitor |
| Alpha-2-macroglobulin | A2MG | P01023 |  | T588 |  | Protease inhibitor |
| Apolipoprotein A-I | APOA1 | P02647 | T103 |  |  | Cholesterol efflux |
| Ceruloplasmin | CERU | P00450 | T1043 | T1052 |  | Iron transport across the cell membrane |
| Clusterin | CLUS | P10909 |  | T202 |  | Apolipoprotein J, clearance of cellular debris and apoptosis |
| Complement C3 | CO3 | P01024 |  | T824 |  | Complement pathway, immune response |
| Complement C4-B | CO4B | P0C0L5 |  | T1124 |  | Complement pathway, immune response |
| Fibrinogen alpha chain | FIBA | P02671 |  | T522; T557 | T374, T412; T557 | Blood clotting |
| Haptoglobin | HPT | P00738 | T119 | T26 |  | Combines with free plasma hemoglobin to allow hepatic recycling of heme iron and to prevent kidney damage |
| Immunoglobulin heavy constant alpha 2 (Fragment) | IGHA2 | A0A0G2JMB2 |  | T71 |  | Antibody |
| Immunoglobulin heavy constant mu | IGHM | P01871 | T82 |  |  | Antibody |
