## Supplemental Figures for "Protein Posttranslational Signatures Identified in COVID-19 Patient Plasma"

**SUPPLEMENTAL ONLINE INFORMATION**

**Vedula_et_al_Fig.S1**

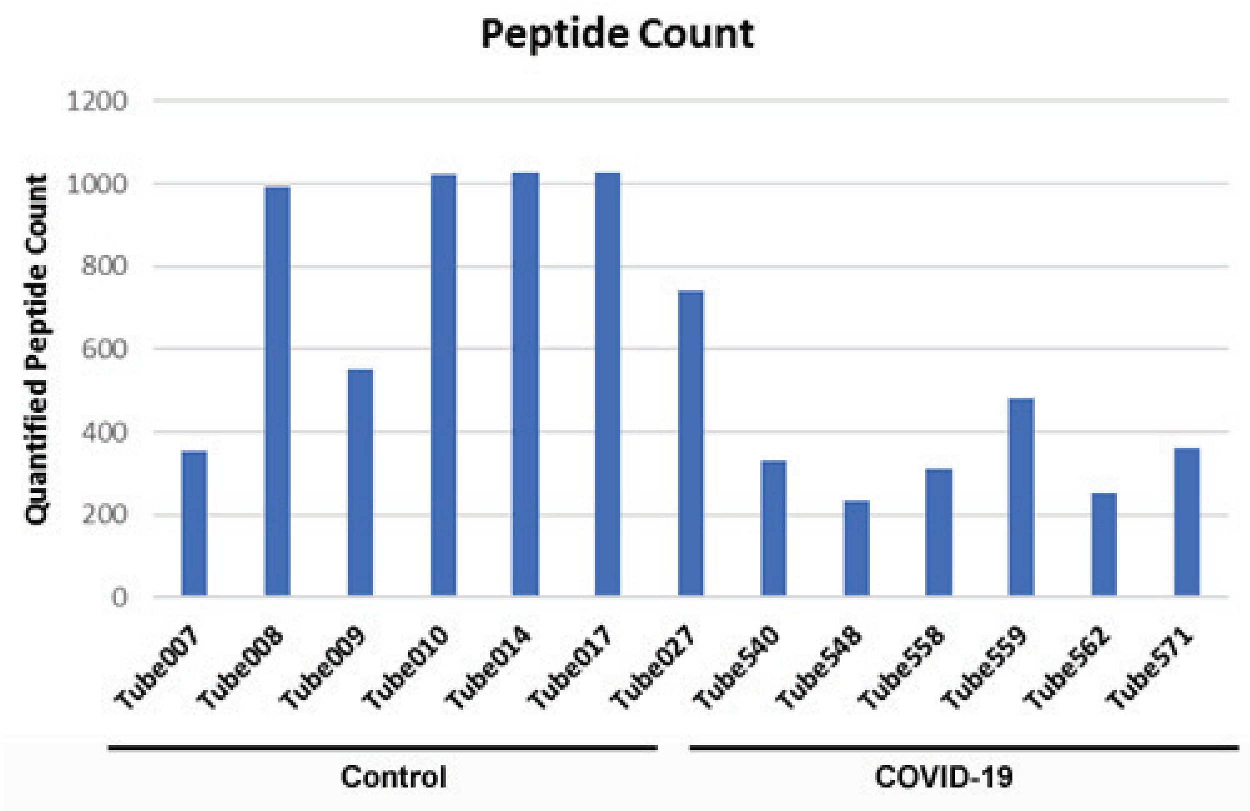


**Figure S1. Overall number of plasma peptides identified by peptidomics is lower in COVID-19 compared to control.** Numbers of identified peptides from each sample are shown, samples 007-027 are control and 540-571 are COVID-19.

**Vedula_et_al_FigS2**

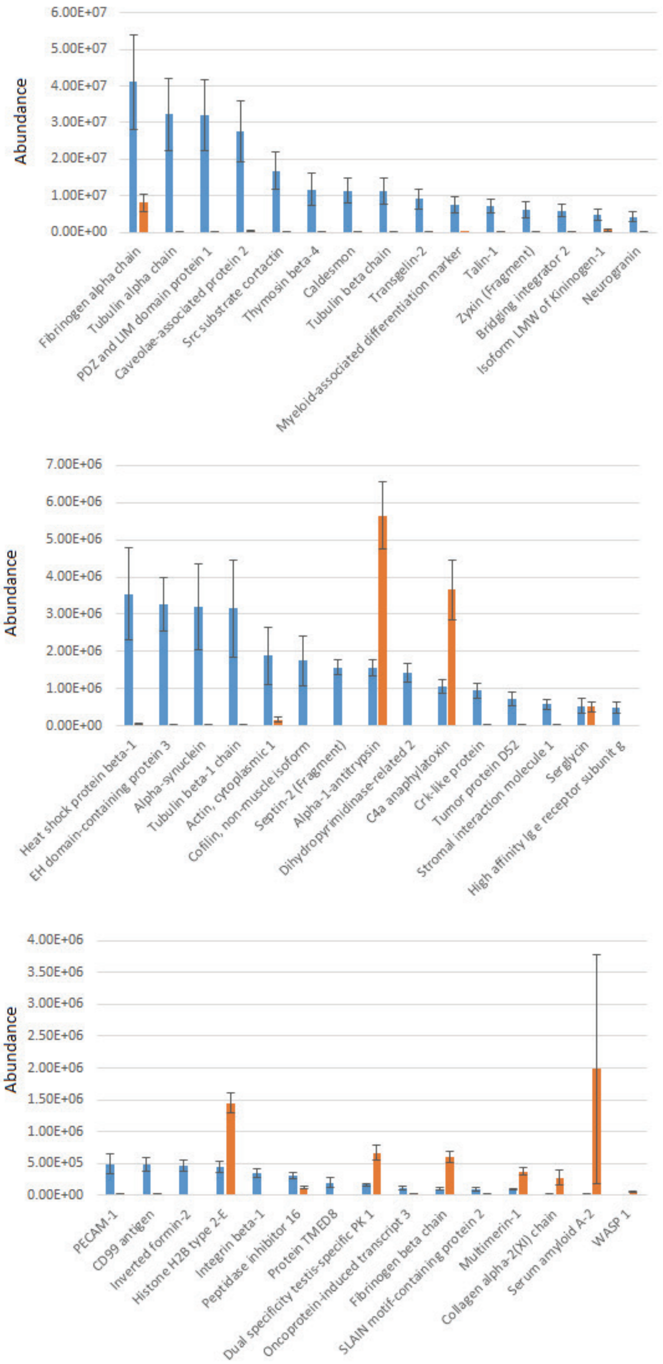


**Figure S2. Plasma peptides from COVID-19 patients exhibits prominent changes in composition compared to control. Combined normalized intensities of all significantly changed peptides for each parent protein listed on the X axis. Bars represent normalized intensity levels averaged for all samples in each group. Proteins with lower combined intensities are shown. See Fig. 1 for the highest abundance hits. Error bars represent SEM (n=7 for control, 6 for COVID-19).**

**Vedula_et_al_Fig.S3**

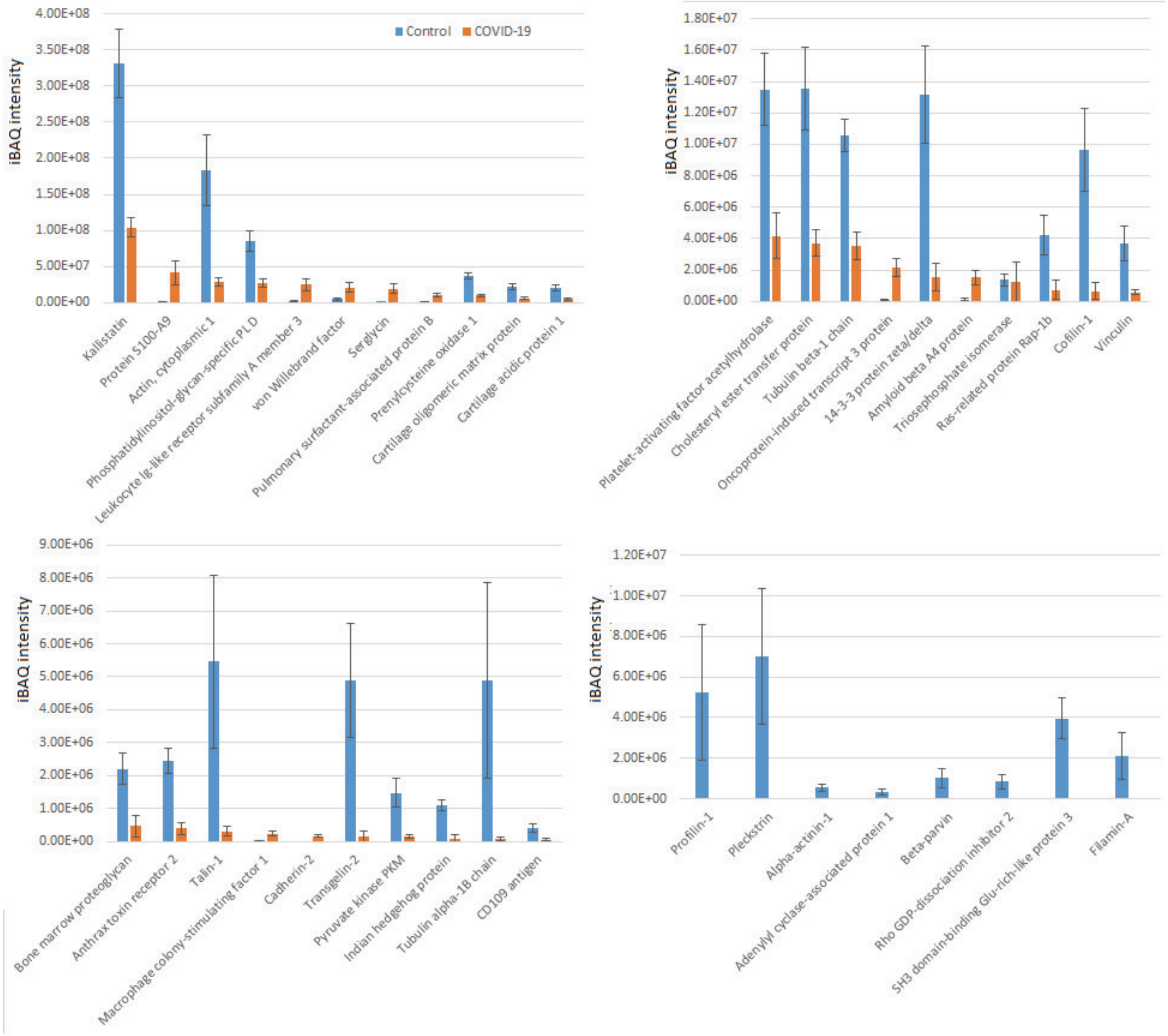


**Figure S3. Plasma proteins from COVID-19 patients exhibits prominent changes compared to control. iBAQ Intensities of the less abundant proteins showing significant differences between COVID-19 and control plotted on different scales. See Fig. 3 for the most abundant hits. Bars represent normalized intensity levels averaged for all samples in each group, error bars represent SEM (n=7 for control, 6 for COVID-19).**

**Vedula_et_al_Fig.S4**

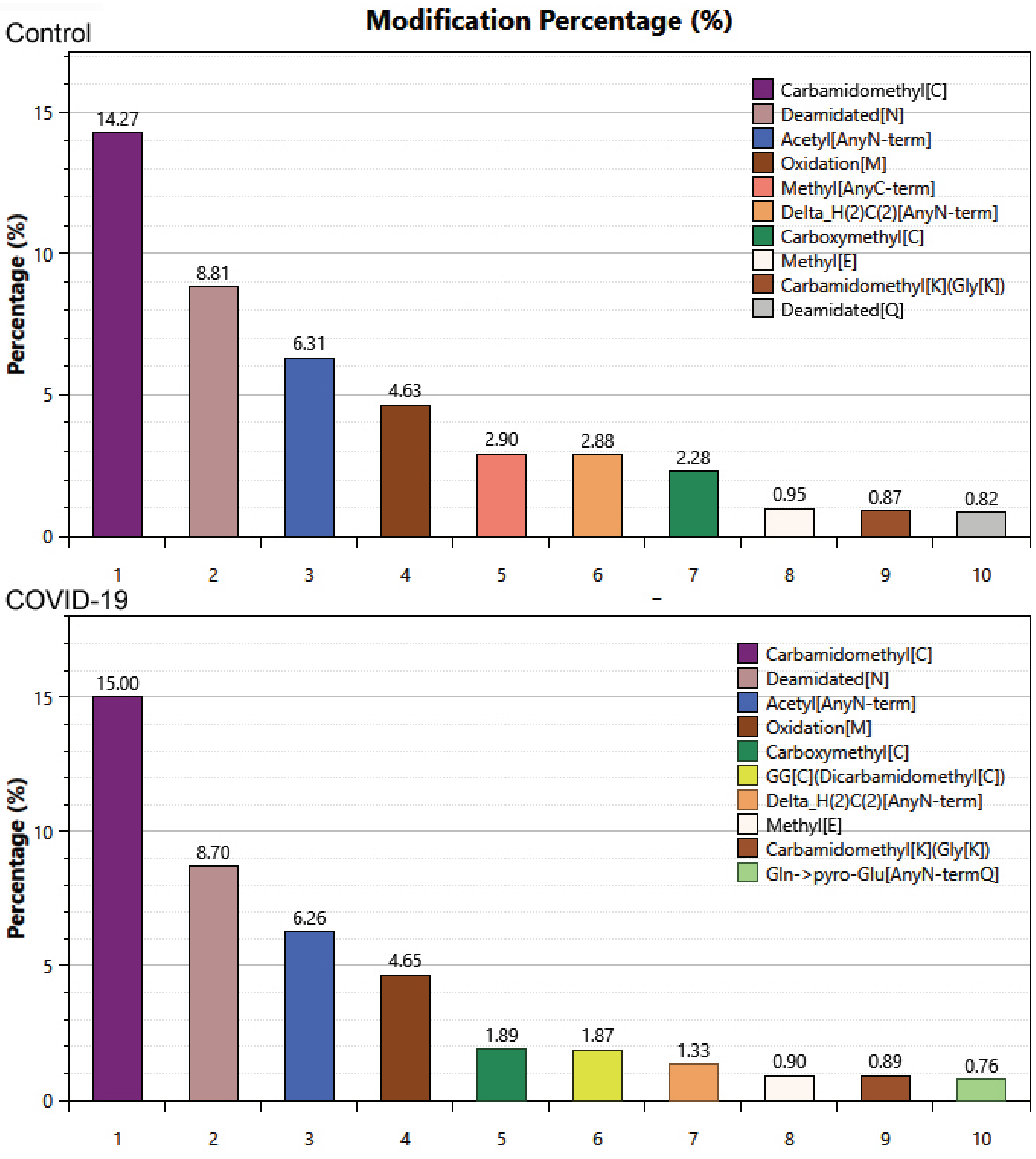


**Figure S4. Top abundance posttranslational modifications identified in control and COVID-19 plasma samples. Percentages of modification derived directly from pFind search are shown.**

**Vedula_et_al_Fig.S5**

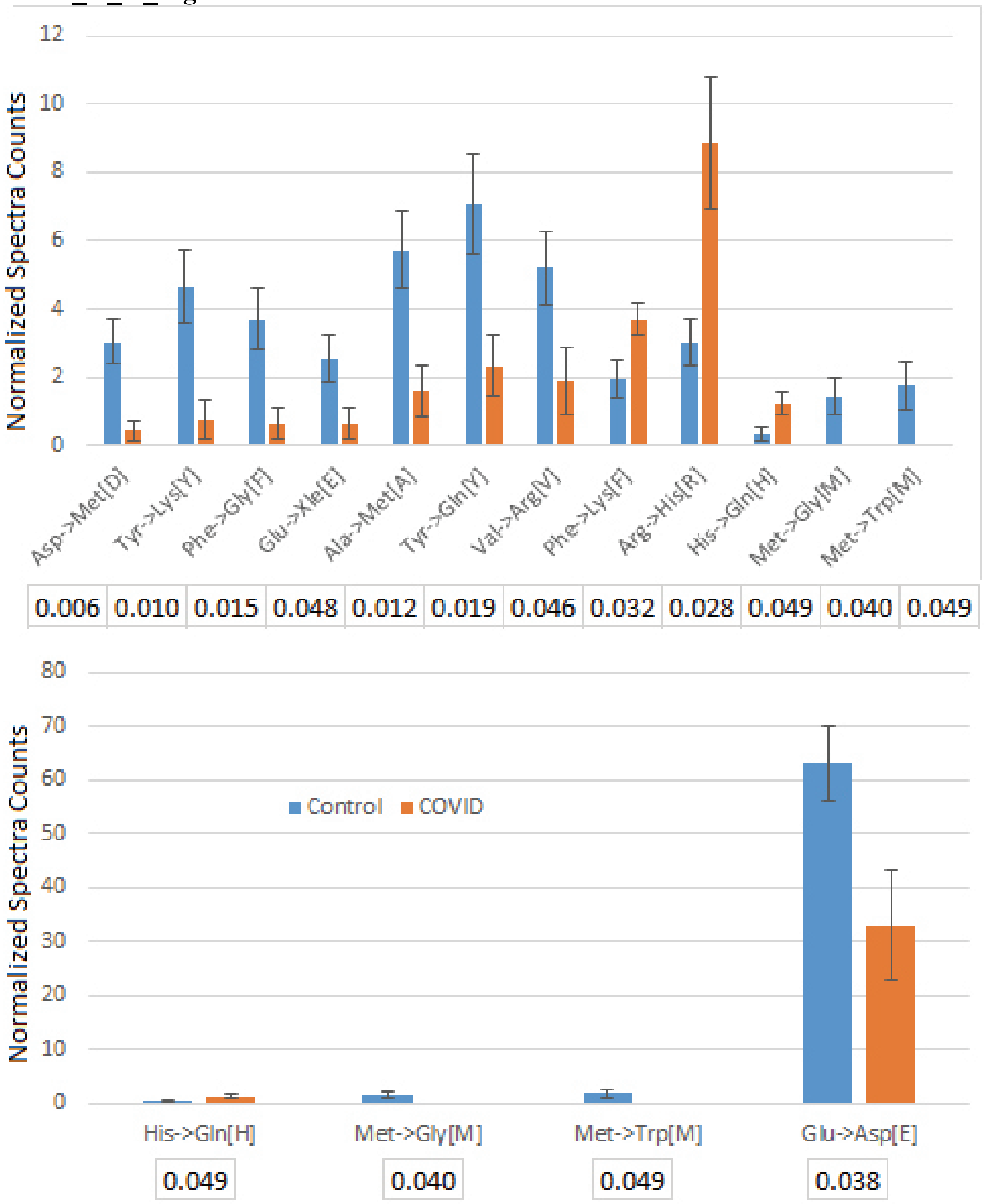


**Figure S5. Plasma proteins from COVID-19 patients exhibits altered patterns of amino acid substitutions indicative of single nucleotide polymorphisms. Error bars represent SEM (n=7 for control, 6 for COVID-19). P values were calculated by 2-tailed Student's T-test. P values calculated by 2-tailed Student's T-test are listed underneath each set. Three last sets on the top chart are also duplicated in the bottom chart for scale.**

**Dataset, Tables and Supplemental Tables:**

**Dataset 1. Sample spectra from peptides identified as arginylated in our analysis.**
**Table S1. List of patient samples used in this study.**

**Table S2. Peptides exhibiting significant differences between control and COVID-19 plasma.**

**Table S3. Peptides from the peptidome analysis that did not exhibit significant differences between control and COVID-19 plasma.**

**Table S4. Proteins exhibiting significant differences between control and COVID-19 plasma.**

**Table S5. Proteins that did not exhibit significant differences between control and COVID-19 plasma.**

**Table S6. Significant PTMs identified by pFind search used for the chart in this study.**

**Table S6. Individual and combined results from pFind search listing the modification sites in each sample.**

**Table S7. Proteins and sites identified as arginylated in control and COVID-19 plasma.**

**Table S8. Proteins and sites identified as phosphorylated on Thr in control and COVID-19 plasma.**
